# Comparative genomics of the major parasitic worms

**DOI:** 10.1101/236539

**Authors:** International Helminth Genomes Consortium

## Abstract

Parasitic nematodes (roundworms) and platyhelminths (flatworms) cause debilitating chronic infections of humans and animals, decimate crop production and are a major impediment to socioeconomic development. Here we compare the genomes of 81 nematode and platyhelminth species, including those of 76 parasites. From 1.4 million genes, we identify gene family births and hundreds of large expanded gene families at key nodes in the phylogeny that are relevant to parasitism. Examples include gene families that modulate host immune responses, enable parasite migration though host tissues or allow the parasite to feed. We use a wide-ranging *in silico* screen to identify and prioritise new potential drug targets and compounds for testing. We also uncover lineage-specific differences in core metabolism and in protein families historically targeted for drug development. This is the broadest comparative study to date of the genomes of parasitic and non-parasitic worms. It provides a transformative new resource for the research community to understand and combat the diseases that parasitic worms cause.

## Introduction

Over a quarter of the human population is infected with parasitic worms, encompassing many species of nematodes (roundworms) and platyhelminths (flatworms)^1^. Although rarely lethal, infections are typified by long-term chronicity, leading to pain, malnutrition, physical disabilities, retarded development, social stigma associated with deformity, and detrimental effects on family members caring for the afflicted. These diseases of poverty impede economic development and attract little medical research investment. Infections with nematodes and platyhelminths make up eight of the WHO list of 20 most neglected tropical diseases of humans and cost the livestock industry billions of dollars annually^2^ through lost production^3^. Nematodes also decimate crop production^4^.

A limited spectrum of drugs are available to treat nematode or platyhelminth infections. Resistance is widespread in farm animals^5^ and mass drug administration in human populations, using repeated monotherapies, is increasing the risk of resistance to human anthelmintics ^6^. Ivermectin, the discovery of which led to a Nobel prize, is used to treat many nematode infections^7^ but like other anthelmintics does not prevent re-infection^8^. There are no vaccines for humans, and very few for animals^9^. The commonly-used plant-parasitic nematicides are environmentally toxic^10^, and need urgent replacement.

In nematodes (phylum Nematoda), parasitism has arisen multiple times^11^. Nematoda is part of the Ecdysozoa superphylum that also contains Arthropoda and several smaller phyla^12^. Nematodes have been classified into five major clades (numbered I to V), four of which contain human-infective parasites and are analysed here (Fig. 1). In platyhelminths (phylum Platyhelminthes), nearly all parasitic groups had a single origin^13^. The majority of parasites are cestodes (tapeworms) and trematodes (flukes) in the class Neodermata (Fig. 1). Platyhelminthes is part of the superphylum Lophotrochozoa along with molluscs, annelids and other groups. Comparing the genome biology of parasitism between these two phyla may reveal common strategies employed to subvert host defences and drive disease processes.

**Fig. 1:**
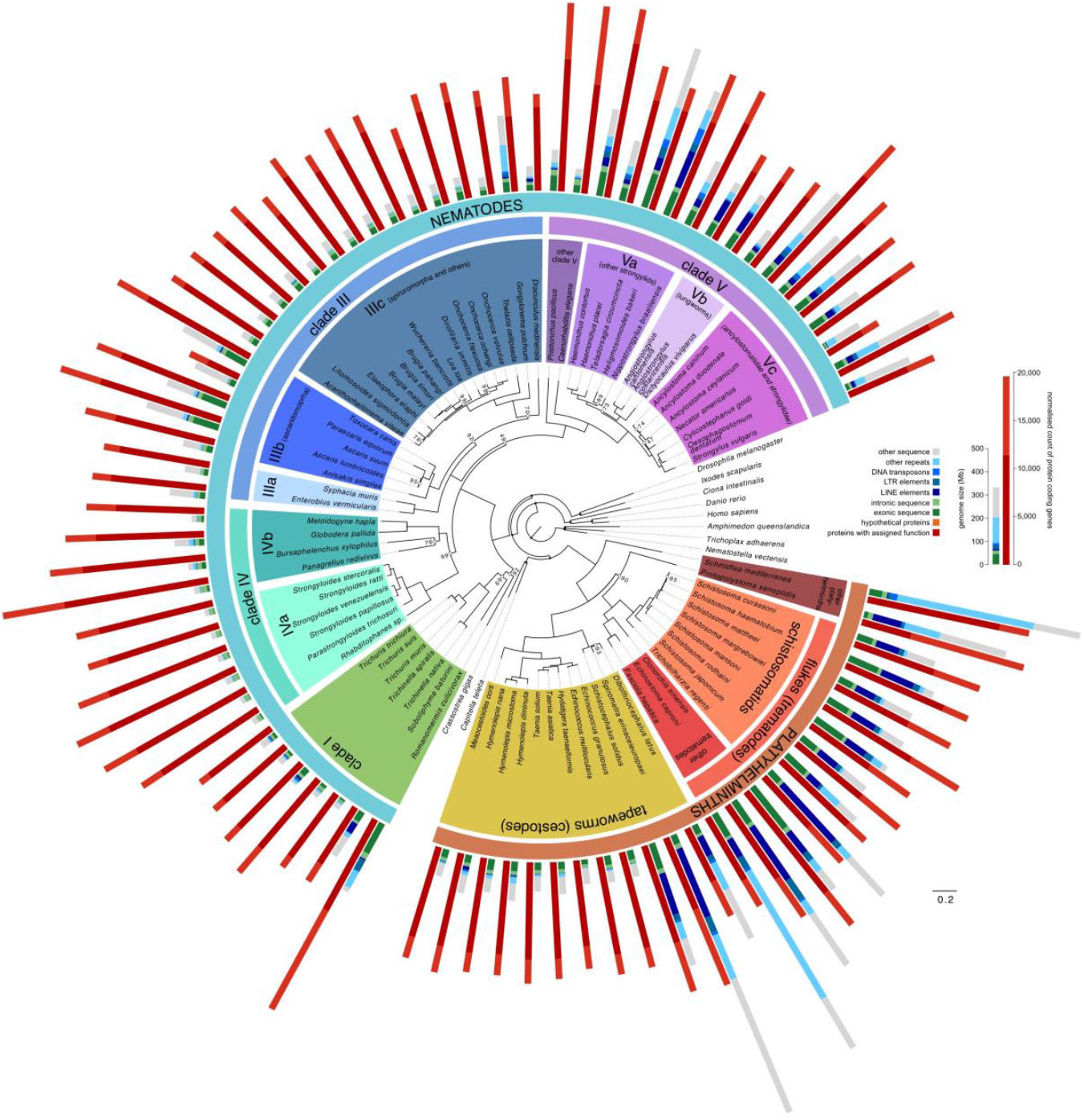
Genome-wide phylogeny of 81 nematode and platyhelminth species and 10 outgroup species. Maximum-likelihood phylogeny based on a partitioned analysis of a concatenated data matrix of 21,649 amino acid sites from 202 single-copy orthologous proteins present in at least 23 of the species. Values on marked nodes are bootstrap support values; all unmarked nodes were supported by 100 bootstrap replicates; nodes with solid marks were constrained in the analysis. Bar plots show genome sizes and total lengths of different genome features, and normalised gene count (Supplementary Information: Results 1.2) for proteins with inferred functions based on sequence similarity (having an assigned protein name; Supplementary Information: Methods 3), or those without (named ‘hypothetical protein’).

In this study, we have combined 36 published genomes with new assemblies for 31 nematode and 14 platyhelminth species into the largest genome comparison to date of parasitic and non-parasitic worms. We have used these data to identify gene families associated with the evolution of major parasitic groups. To accelerate the search for new interventions, we have mined the dataset of more than 1.4 million genes to predict new drug targets and drugs.

## Results

### Genomic diversity in parasitic nematodes and platyhelminths

We have produced draft genomes for 45 nematode and platyhelminth species (Supplementary Table 1, Extended Data Fig. 1) and predicted 0.8 million new protein-coding genes, with 9, 132–17,274 genes per species (5–95% percentile range; Supplementary Table 2, Extended Data Fig. 1, Supplementary Information: Results 1.1, 1.2). We combined these new data with 36 previously published worm genomes, comprising one free-living and ten parasitic platyhelminths, four free-living (including the model *Caenorhabditis elegans)* and 21 parasitic nematodes, and 10 non-parasitic outgroup taxa from other animal phyla (Supplementary Tables 3, 4 and Fig. 1), into a comparative genomics resource of 91 species (Fig. 1).

Genome size varied greatly within each phylum, from 42–700 Mb in nematodes, and 104–1,259 Mb in platyhelminths. Repeat content of the genomes also ranged widely, from 3.8–54.5% (5–95% percentile; Supplementary Table 5). To rank factors driving genome size variation in these species, we built a multiple regression model and identified long terminal repeat transposons, simple repeats, assembly quality, DNA transposable elements, total length of introns, and low complexity sequence as being the most important (Supplementary Information: Results 1.3, Supplementary Table 6). Genome size variation is thus largely due to changes in different non-coding elements, as expected^14^, suggesting it is either non-adaptive or responding to selection only at the level of overall genome size.

### Novel gene families and gene family evolution

We inferred gene families from the predicted proteomes of the 91 species using Ensembl Compara (which identifies families based on all-versus-all BLASTP comparisons^15^). Of the 1.6 million proteins, 1.4 million were placed into 108,351 families, for which phylogenetic trees were built and orthology and paralogy inferred (Supplementary Information: Methods 8, Supplementary Fig. 1, Supplementary Table 7). Species trees inferred from 202 single-copy gene families that were present in at least 25% of species (Fig. 1), or from presence/absence of gene families, largely agreed with the expected species and clade relationships, except for a couple of known contentious issues (Extended Data Fig. 2).

Gene families specific to, or with significantly changed membership in, particular parasite clades compared to free-living or host taxa are likely to reflect important changes in parasite biology and may be targeted by new antiparasitic interventions (Supplementary Fig. 1). Parasitic species contained significant novelty in gene content. For example, ~28,000 parasitic nematode gene families contained members from two or more parasitic species but were absent from *C. elegans*. We identified 5881 clade-specific families (synapomorphies) at key nodes in the phylogeny relevant to parasitism (Supplementary Information: Results 2.2, Supplementary Table 8). These were either gene family births, or subfamilies that were so diverged from their homologues that they were identified as separate families. Functional annotation of these synapomorphic families was diverse (Fig. 2) but they were frequently associated with sensory perception (such as G-protein coupled receptors; GPCRs), parasite surfaces (platyhelminth tegument or nematode cuticle maintenance proteins), and protein degradation (proteases and protease inhibitors).

**Fig. 2:**
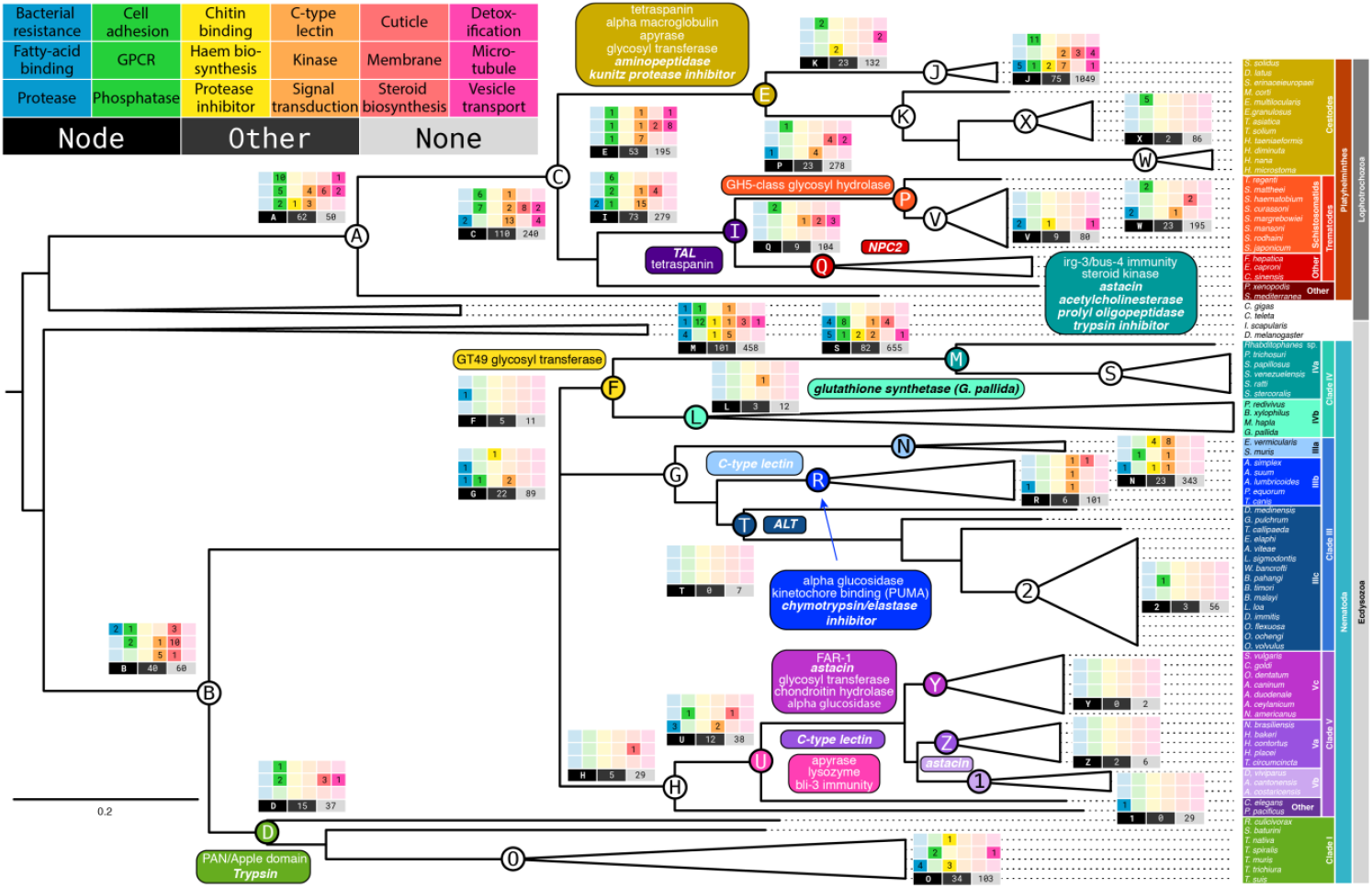
Functional annotation of synapomorphic and expanded gene families. Rectangular matrices indicate counts of synapomorphic families grouped by 18 functional categories, detailed in the major panel. Representative functional annotation of a family was inferred if more than 90% of the species present contained at least one gene with a particular domain. The node in the tree to which a panel refers is indicated in each matrix. ‘Other’ indicates families with functional annotation that could not be grouped into one of the 18 categories. ‘None’ indicates families that had no representative functional annotation. Curved panels indicate gene families showing noteworthy expansions at nodes. Previously published expansions are in bold Italics.

Amongst nematodes, clade IVa (which includes *Strongyloides* spp, Fig. 1) displayed the highest number of clade-specific gene families, including a novel ferrochelatase-like (FeCL) family. Most nematodes lack functional ferrochelatases for the last step of haem biosynthesis^16^, but harbour FeCL genes of unknown function, to which the synapomorphic clade IVa family was similar (Supplementary Fig. 2). Exceptions are animal parasites in nematode clades III (e.g. the ascarids and the filaria) and IV that acquired a functional ferrochelatase via horizontal gene transfer^17,18^.

Within the parasitic platyhelminths (Neodermata) that includes *Protopolystoma*, trematodes and cestodes, a clade-specific inositol-pentakisphosphate 2-kinase (IP2K) was identified. In some species of *Echinococcus* tapeworms IP2K produces inositol hexakisphosphate nano-deposits in the extracellular wall (the laminated layer) that protects larval metacestodes^19^. The deposits increase the surface area for adsorption of host proteins and may promote interactions with the host^20^.

We identified 244 gene family expansions in parasites (Supplementary Table 9a; Supplementary Information: Results 2.3, of which 32 had no functional annotation. Of those with some level of functional information, 25 corresponded to previously described expansions, including 21 associated with biological functions important for parasitism (Fig. 2). A further 43 were placed into major functional classes that historically have been most successful as drug targets (kinases, GPCRs, ion channels and proteases^21^; Supplementary Table 9a).

Among the remaining 144 newly identified expansions were families that interact with host immune systems (Fig. 2). *Taenia* tapeworms and clade V strongylid nematodes (i.e. Va, Vb, and Vc; Fig. 1) contained two expanded families with apyrase domains that may have a role in hydrolysing ATP (a host danger signal) from damaged host tissue^22^ (Extended Data Fig. 3a-b). Moreover, many of the strongylid members also contained amine oxidoreductase domains, possibly to reduce production of pro-inflammatory amines, such as histamine, from host tissues^23^. In platyhelminths, we observed expansions of tetraspanin families that are likely components of the host-pathogen interface. Examples have been described that are part of extracellular vesicles released by helminths within their hosts^24^; or that bind the Fc domain of host antibodies^25^; or that are highly immunogenic^26^ (Extended Data Fig. 3c-d). In strongylids, especially clade Vc, an expansion of the fatty acid and retinol-binding (FAR) gene family, implicated in host-parasite interaction in plant- and animal-parasitic nematodes^27,28^ (Extended Data Fig. 3e), suggests a role in immune modulation. Repertoires of glycosyl transferases have expanded in nematode clades Vc and IV, and tapeworms (Extended Data Fig. 4a-c) and may be used to evade or divert host immunity by modifying parasite surface molecules directly exposed to the immune system^29^; alternatively, the surface glycoprotein may interact with lectin receptors on innate immune cells in an inhibitory manner ^30^. An expanded chondroitin hydrolase family in nematode clade Vc may possibly be used either for larval migration through host connective tissue or to digest host intestinal walls (Extended Data Fig. 3f). Similarly, a GH5 glycosyl hydrolase family expanded in schistosomes contained genes with egg-enriched expression^31,32^, and may be used by eggs to traverse host tissues such as bladder or intestinal walls (Extended Data Fig. 3g). In nematode clade I, we found an expansion of a family with the PAN/Apple domain, which is implicated in attachment of some protozoan parasites to host cells^33^, and possibly modulates host lectin-based immune activation (Extended Data Fig. 3h).

We identified several expansions of gene families involved in innate immunity of the parasites, as well as their development. These included families implicated in protection against bacterial or fungal infections in nematode clade IVa *(bus-4* GT31 galactosyltransferase^34^, *irg-3*^35^) and clades Va/Vc (lysozyme^36^ and the dual oxidase *bli-3*^37^) (Supplementary Fig. 3a-d). In nematode clade IIIb, a family was expanded that contains orthologs of the *Parascaris* coiled-coiled protein PUMA, involved in kinetochore biology^38^ (Supplementary Fig. 3e). This expansion may be possibly related to the evolution in this group of chromatin diminution, which results in an increase in the number of chromosomes requiring correct segregation during metaphase^39^. In nematode clade IVa and in *Bursaphelenchus*, an expansion of a steroid kinase family (Supplementary Fig. 3f) is suggestive of novelty in steroid-regulated processes in this group, such as the switch between free-living or parasitic stages in *Strongyloides* ^40^.

The SCP/TAPS (Sperm-Coating Protein/Tpx/Antigen 5/Pathogenesis-related protein 1) genes are a particularly striking example of a family associated with parasitism through their abundance, secretion and immunomodulatory properties^41^ but whose functions remain unclear (Supplementary Information: Results 3.1). There were eight expanded families containing SCP/TAPS but a phylogenetic analysis of the full repertoire of 3,167 sequences revealed intra- and inter-specific expansions and diversification over different evolutionary time-scales (Fig. 3 and Supplementary Fig. 4). In particular, the SCP/TAPS superfamily has expanded independently in nematode clades V (18-381 copies in each species) and in clade IVa parasites (39-166 copies) (Fig. 3, Supplementary Fig. 5, Supplementary Table 10). *Dracunculus medinensis* was unusual in being the only member of clade III to display an expansion (66 copies), which may reflect modulation of the host immune response during the tissue migration phase of the large adult females.

**Fig. 3:**
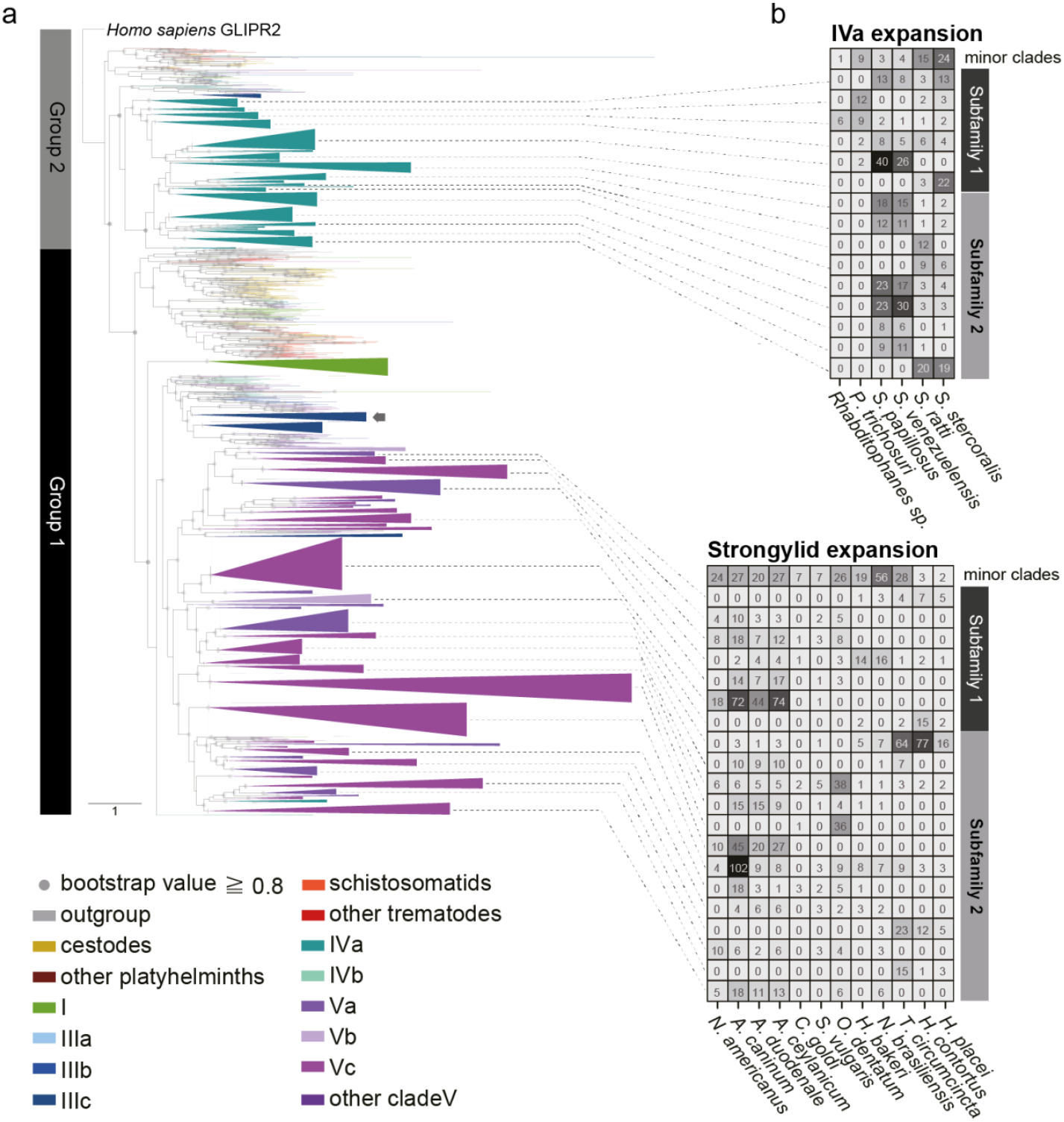
Distribution and phylogeny of SCP/TAPS genes. (a) A maximum-likelihood tree of SCP/TAPS genes. Colours represent different species groups. *H. sapiens* GLIPR2 was used to root the tree. Grey dots show high bootstrap values (≥0.8). A clade was collapsed into a triangle if more than half its leaves were genes from the same species group. Clade I had fewer counts, but was collapsed to show its relationship to other clades’ expansions. A *Dracunculus* expansion is indicated with an arrow. (b) Clades containing more than 15 (in group 1) and 10 (in group 2) genes are shown in a heatmap. ‘Strongylid’ refers to clades Va, Vb and Vc.

Half (46.9%) of the gene families lacked functional annotation. These families tended to be smaller than those with annotations (Supplementary Fig. 6) and in many cases, may correspond to families that are so highly diverged that ancestry cannot be traced, but may also reflect the huge breadth of unexplored parasite biology. Family 393312, for example, contained 96 members, 26% of which were predicted to be secreted, and was highly expanded in clade Va nematodes (Supplementary Fig. 7; Supplementary Table 9a). Even when families could be functionally annotated to some extent (e.g. based on a protein domain), discerning their precise biological role was in many cases a challenge. For example, a sulfotransferase gene family that was expanded in flukes compared with tapeworms included the S. *mansoni* locus *Smp_089320* that is implicated in resistance to the drug oxamniquine^42^ but the endogenous substrate for this enzyme is unknown (Supplementary Fig. 7j).

### Proteins historically targeted for drug development

GPCRs, kinases, proteases and ion channels dominate the list of targets for existing drugs for human diseases^21^, and are attractive leads for development of new ones. We therefore explored the diversity of these key superfamilies across the nematodes and platyhelminths (Fig. 4; Supplementary Information: Results 3.2, 3.3, 3.4, 3.6).

**Fig. 4:**
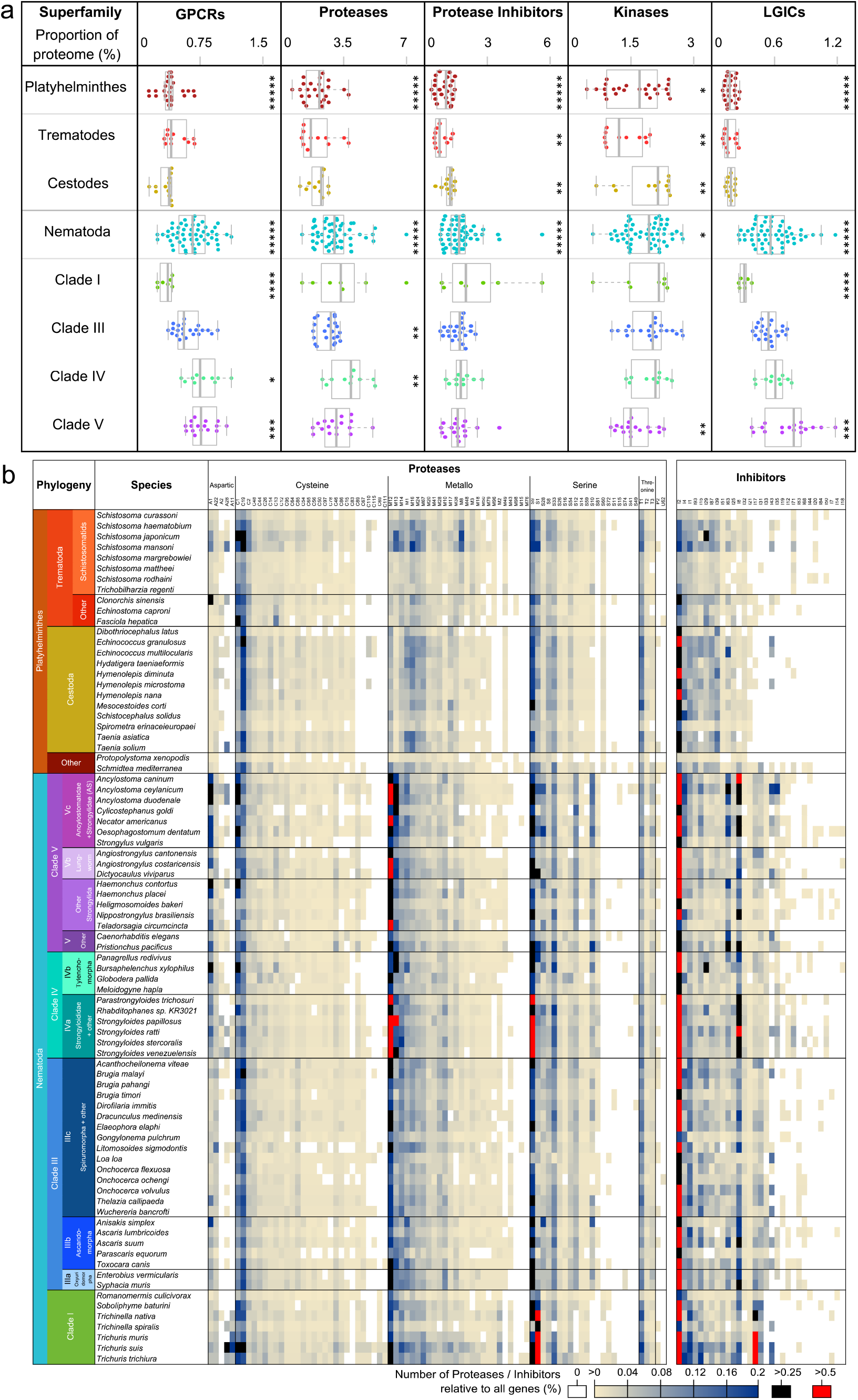
Abundances of superfamilies historically targeted for drug development. (a) Abundances (relative to the total numbers of predicted genes) of genes belonging to superfamilies of interest. Comparisons are (i) platyhelminths vs nematodes, (ii) trematodes vs cestodes and (iii) nematode clade vs other nematodes. * *P*<0.05, ** *P*<0.01, *** *P*<10^−3^, **** *P*<10^−4^, ***** *P*<10^−5^ (Mann Whitney U test). The *C. elegans* GPCR value of 4.6% was not shown. (b) Relative abundance profiles for 84 protease and 31 protease inhibitor families represented in at least 3 of the 81 nematode and platyhelminth species. 33 protease families and 6 protease inhibitor families present in fewer than 3 species were omitted from the visualisation.

Proteases and protease inhibitors perform a number of different functions in parasites, including immunomodulation, host tissue penetration, modification of the host environment (e.g. anticoagulation), and digestion of blood^43^. M12 astacins had particularly expanded in nematode clade IVa (five family expansions), as previously reported^44^, but also in clades Vc and Vb (two additional expansions; Fig. 4, Supplementary Fig. 8, Supplementary Table 11). As many of these species invade through skin (IVa, Vc; Supplementary Table 12) and migrate through the digestive system and lung (IVa, Vc, Vb; Supplementary Table 13), this was consistent with evidence that astacins are involved in skin penetration and migration through connective tissue^45^. The cathepsin B C1-cysteine proteases had particularly expanded in species that feed on blood (two expansions in nematode clades Vc and Va^46^, with highest playthelminth gene counts in schistosomatids and the liver fluke *Fasciola*^47^; Supplementary Fig. 8). Indeed, they are involved in blood digestion in both adult nematodes^48^ and playhelminths^49^, but some likely have different roles such as larval development^50^ and host invasion^51^.

Different families of protease inhibitors (PI) may modulate the activity of parasite proteases or protect parasitic nematodes and platyhelminths from degradation by host proteases, facilitate feeding or manipulate the host response to the parasite^52^. The I2 (Kunitz-BPTI) trypsin inhibitors were the most abundant PIs across all parasitic nematodes and platyhelminths (Fig. 4b). An expansion of the I17 family, which includes secretory leukocyte peptidase inhibitor, was reported previously in *Trichuris muris^53^* but the striking confinement of this expansion to most of the parasites of clade I is now apparent (Fig. 4b). We also observed a notable family of α-2-macroglobulin (I39) PIs that were present in all platyhelminths but expanded in tapeworms (Supplementary Fig. 8). The tapeworm α-2-macroglobulins may be involved in reducing blood clotting at attachment or feeding sites; alternatively, they may reduce activity of defensive signalling proteins from the host, since α-2-macroglobulins bind several cytokines and hormones^54^. Chymotrypsin/elastase inhibitors (family I8) were particularly expanded in clades Vc and IVa (consistent with upregulation of I8 genes in *Strongyloides* parasitic stages^44^) and to a lesser extent in clade IIIb (Fig. 4b), consistent with evidence that these inhibitors may protect *Ascaris* from host proteases^55^. In our analyses we also identified numerous protein domain combinations that were specific to either nematodes or platyhelminths (131 and 50 domain combinations, respectively). Many of these protein innovations involved protease and PI domains. In nematodes, several combinations included Kunitz PI domains, and in platyhelminths metalloprotease families M18 and M28 were found in novel combinations (Supplementary Table 14; Supplementary Information: Results 3.7).

Of the 230 gene families annotated as GPCRs (Fig. 4a and Extended Data Fig. 5), only 21 were conserved across phyla. Chemosensory GPCRs, while abundant in nematodes, were not identified in platyhelminths, although they are identifiable in other Lophotrochozoa (such as Mollusca^56^), suggesting that either the platyhelminths have lost this class or they are very divergent (Supplementary Table 15). GPCR families lacking sequence similarity with known receptors included the platyhelminth-specific rhodopsin-like orphan families (PROFs), which are likely to be Class A receptors and peptide responsive, and several other fluke-specific non-PROF GPCR families. The massive radiation of chemoreceptors in *C. elegans* was unmatched in any other nematode species (87% vs ≤48% of GPCRs). All parasitic nematodes possessed chemoreceptors, with the most in clade IVa, including several large families synapomorphic to this clade (Extended Data Fig. 5), perhaps related to their unusual life cycles that alternate between free-living and parasitic forms.

Our analysis of pentameric ligand gated ion channels (pLGICs) suggested that the ancestor of each phylum lacked a complete complement of pLGIC subunit classes (Supplementary Fig. 9, Supplementary Table 16). Divergence between nematode and platyhelminth pLGICs resulted from independent expansion and functional divergence. For example, glutamate signalling arose independently in platyhelminths and nematodes^57^, and in trematodes the normal role of acetylcholine has been reversed, from activating to inhibitory^58^. Our analysis suggested the platyhelminth acetylcholine-gated anion channels are most related to the Acr-26/27 group of nematode nicotinic acetylcholine receptors that are the target of the anthelmintics morantel and pyrantel^59^, rather than to nematode acetylcholine-gated cation channels, targeted by nicotine and levamisole (Supplementary Fig. 10).

ABC transporters (Supplementary Table 17) showed similar loss and independent expansion between nematodes and platyhelminths. The P-glycoprotein class, responsible for the transport of environmental toxins and linked with anthelmintic resistance, was expanded relative to vertebrates^60^, with increased numbers in nematodes (Supplementary Fig. 11).

### Metabolic reconstructions of nematode and platyhelminth parasites

We generated metabolic reconstructions for each of the 81 nematode and platyhelminth species, based on high confidence assignment of enzyme classes (Supplementary Table 18a). The nematodes in our dataset clearly had a greater range of annotated enzymes per species than did the platyhelminths (Extended Data Fig. 6a), in part reflecting the paucity of biochemical studies in platyhelminths. Because variation in assembly quality or divergence from model organisms could bias these inferences, we identified losses of pathways and differences in pathway coverage across different clades (Supplementary Information: Results 4, Fig. 5, Supplementary Fig. 12). Pathways related to almost all metabolic superpathways in KEGG^61^ showed significantly lower coverage for platyhelminths (versus nematodes) and filaria (versus other nematodes) (Extended Data Fig. 6b).

In contrast to most animals, nematodes possess the glyoxylate cycle, which enables conversion of lipids to carbohydrates that can be used for biosyntheses (e.g. during early development), and to avert starvation^62^. The glyoxylate cycle appeared to have been lost independently in the filaria and *Trichinella* species (Fig. 5a; M00012), both of which are tissue-dwelling obligate parasites. The filaria and *Trichinella* had also independently lost alanine-glyoxylate transaminase, which converts glyoxylate to glycine (Fig. 5b). Glycine can be converted by the glycine cleavage system (GCS) to 5,10-methylenetetrahydrofolate, a useful one-carbon pool for biosyntheses, and two key GCS proteins appeared to have been lost independently from filaria and tapeworms, suggesting their GCS is non-functional (Supplementary Table 19e). Additionally, filaria had lost the ability to produce and use ketone bodies, a useful temporary store of acetyl CoA under starvation conditions (Supplementary Table 19b). *Dracunculus medinensis*, the outgroup to the filaria in clade IIIc, does have a free-living L1 stage (Supplementary Table 12) and had not lost the glyoxylate cycle, GCS, or usage of ketone bodies.

**Fig. 5:**
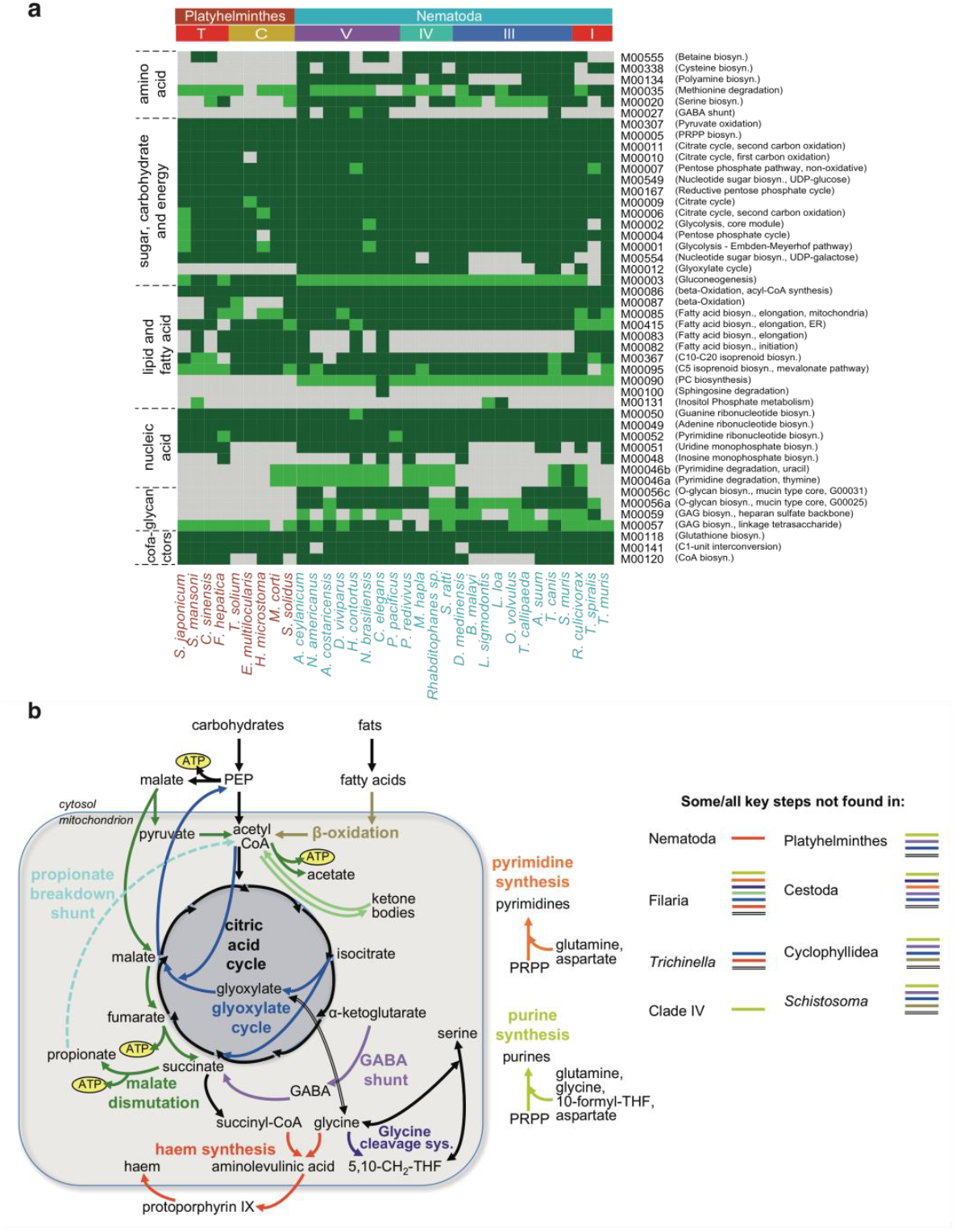
Metabolic modules and biochemical pathways in platyhelminths and nematodes. (a) Topology based detection of KEGG metabolic modules among tier 1 (with high quality assemblies; see Supplementary Information: Methods 7) species (dark green = present; light green = largely present (only one enzyme not found). Only modules detected to be complete in at least one species are shown. The EC annotations used for this figure included those from pathway hole-filling and those based on Compara families (Supplementary Table 18a, b). (b) Biochemical pathways that appear to have been completely or partially lost from certain platyhelminth and nematode clades.

Multiple initial steps of pyrimidine synthesis were absent from some nematodes, including all filaria (as previously reported^63^) and tapeworms, suggesting these taxa obtain pyrimidines from their hosts or from *Wolbachia* endosymbionts, in the case of many filaria (Supplementary Table 19f). Platyhelminths and many nematodes (especially clade IVa and filaria) were also devoid of key enzymes in purine synthesis (Supplementary Table 19g) so likely rely on salvage instead. However, in contrast to the widespread belief that nematodes cannot synthesise purines^64,65^, complete or near-complete purine synthesis pathways were found in most members of clades I, IIIb and V. Nematodes are known to be unable to synthesise haem^16^ but the pathway was present in platyhelminths, including S. *mansoni* (despite conflicting biochemical data^16^ (Supplementary Table 19h; Supplementary Table 20i).

Genes from the β-oxidation pathway, used to break down lipids as an energy source, were absent from schistosomes and some cyclophyllidean tapeworms *(Hymenolepis, Echinococcus;* Fig. 5a, M00087; Supplementary Table 19a). These species live in glucose-rich environments and may have evolved to use glucose and glycogen as principal energy sources. However, biochemical data suggest they do perform β-oxidation^66,67^, so they may have highly diverged but functional β-oxidation genes.

The lactate dehydrogenase (LDH) pathway is a major source of ATP in anaerobic but glucose-rich environments. Platyhelminths had high numbers of *LDH* genes, as did blood-feeding *Ancylostoma* hookworms (Supplementary Fig. 13g). Nematode clades Vc (which includes *Ancylostoma)* and IIIb had expansions of α-glucosidases that can break down starch and disaccharides in host food to glucose (Supplementary Fig. 13a). Many nematodes and flatworms use malate dismutation as an alternative pathway for anaerobic ATP production ^68^. The importance of the pathway for clade IIIb nematodes was reflected in their gene content. We identified expanded families encoding two key pathway enzymes PEPCK and methylmalonyl CoA epimerase, and the intracellular trafficking chaperone for cobalamin (vitamin B-12), a cofactor for the pathway (Supplementary Fig. 13c-e; Supplementary Table 9a). A second cobalamin-related family (CobQ/CbiP) was clade IIIb-specific and appeared to have been gained from bacteria (Supplementary Fig. 14a). A glutamate dehydrogenase family expanded in clade IIIb (Supplementary Fig. 13h) is consistent with a GABA shunt that helps maintain redox balance during malate dismutation. In clade Va we observed expansion of a gene family belonging to a propionate breakdown pathway ^69^ (Supplementary Fig. 13f), which may degrade propionate from malate dismutation or fermentation in the host’s stomach^70^. Clade I nematodes had an acetate/succinate transporter that appeared to have been gained from bacteria, and is likely to participate in acetate/succinate uptake or efflux (Supplementary Fig. 14b).

### Identifying new anthelmintic drug targets and drugs

Many of the 24 WHO-listed anthelmintic drugs are compromised by low efficacy, side-effects or rising resistance in parasite populations. To reveal potential new drug targets and drugs, we developed a pipeline to identify the most promising targets from parasitic nematodes and platyhelminths, and compounds likely to interact with them from the ChEMBL database (Extended Data Fig. 7; Supplementary Information: Results 5). For the top 15% of highest-scoring targets (n=289), including 17 out of 19 known or likely targets for WHO-listed anthelmintics that are represented in ChEMBL, the pipeline identified compounds that are predicted to interact (Supplementary Table 21b). This potential screening set, when collapsed by chemical class, contained 5046 drug-like compounds, including 817 drugs with phase III or IV approval and 4229 medicinal chemistry compounds (Supplementary Table 21d). We used a self-organising map (SOM) to cluster these compounds based on their molecular fingerprints (Fig. 6a). This classification showed that the screening set was significantly more structurally diverse than existing anthelmintic compounds (Fig. 6b).

**Fig. 6:**
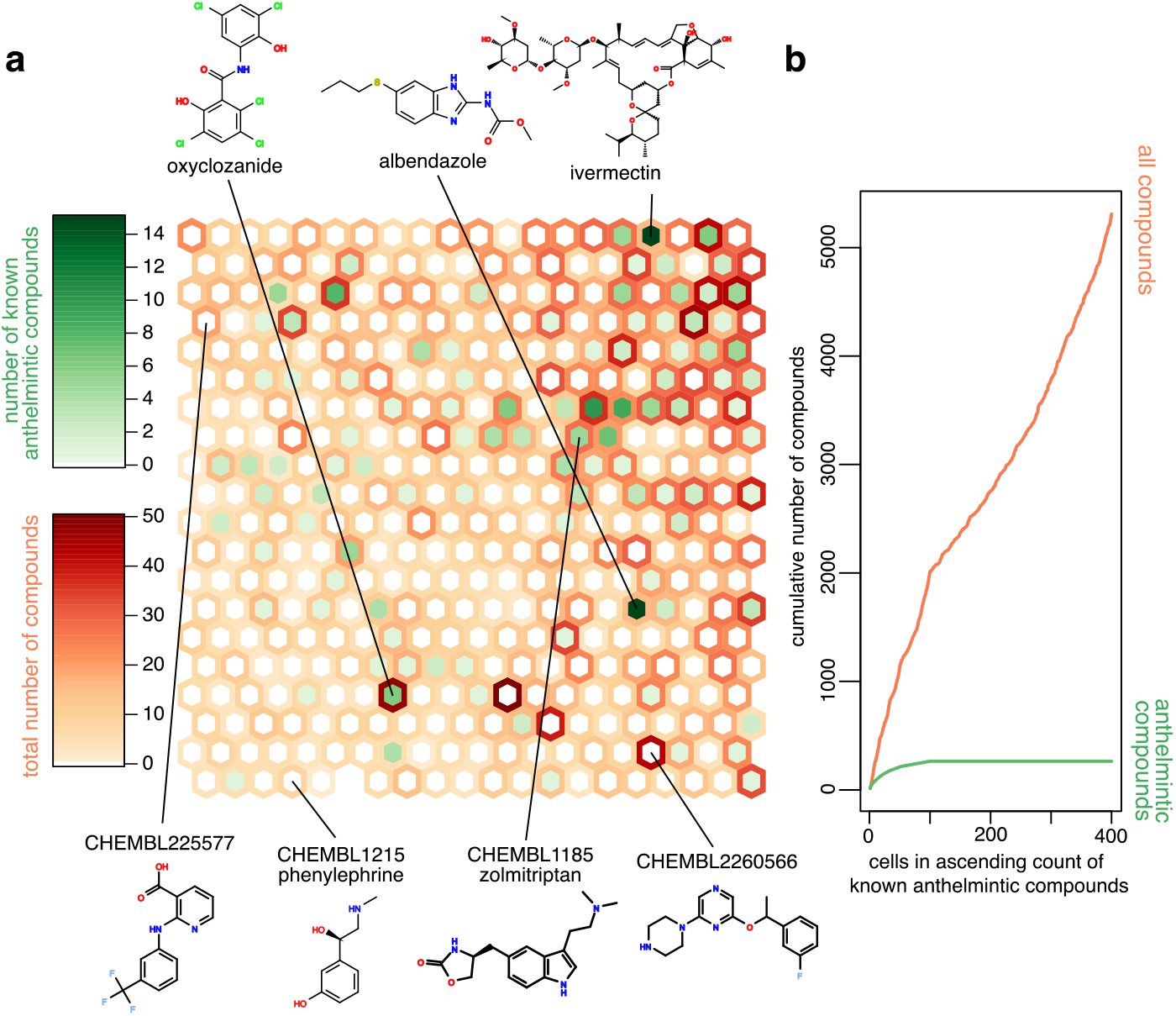
Self-organising map of known anthelmintic compounds and our screening set of 5046 drug-like compounds. (a) A self-organising map (SOM) clustering known anthelmintic compounds (Supplementary Table 21a) and our screening set of 5046 compounds. The density of red and green shows the number of screening set and known anthelmintic compounds clustered in each cell, respectively. Structures for representative known anthelmintic compounds are shown at the top, and for screening set compounds along the bottom. (b) Cumulative counts of known anthelmintic and screening set compounds, showing that known anthelmintic compounds are more highly clustered in the SOM. Thus, our screening set represents a more diverse sample of chemical space than the known anthelmintic compounds.

The 289 targets were further prioritised to 40 high-priority targets, based on predicted selectivity, avoidance of side-effects (clade-specific chokepoints or lack of human homologues), and putative vulnerabilities, such as those suggested by gene family expansions in parasite lineages, or belonging to pathways containing known or likely anthelmintic targets (Extended Data Fig. 8). These 40 targets were associated with 720 drug-like compounds comprising 181 phase III/IV drugs and 539 medicinal chemistry compounds. There is independent evidence that some of these have anthelmintic activity. For example, we identified several compounds that potentially target glycogen phosphorylase, which is in the same pathway as a likely anthelmintic target (glycogen phosphorylase phosphatase, likely target of niridazole; Extended Data Fig. 8). These compounds included the phase III drug alvocidib (flavopiridol), which has anthelmintic activity against *C. elegans^71^*. Another example is the target cathepsin B, greatly expanded in nematode clade Va (Supplementary Table 9a), for which we identified several compounds including the phase III drug odanacatib, which has recently been shown to have anthelmintic activity against hookworms^72^. Existing drugs such as these are attractive candidates for repurposing and fast-track therapy development, while the medicinal chemistry compounds provide a starting point for broader anthelmintic screening.

## Discussion

The evolution of parasitism in nematodes and platyhelminths occurred independently, and started from very different ancestral gene sets and physiologies. Despite this, the common selective pressures of adaptation to host gut, blood or tissue environments, the specific nutrient abundances available within hosts, the need to avoid or subvert the host immune system and the acquisition of complex multi-host/environment lifecycles to effect transmission, may have driven adaptations in common biological pathways and therefore common genomic solutions. We have surveyed diverse parasitic worms, with a focus on those infecting humans and livestock. Despite the diversity of the species analysed we identified common themes. Gene family analyses revealed expansions in functional classes that reflect the importance to the parasitic lifestyle. We were able to identify biochemical pathways uniquely absent from, or present in, particular parasite groups. Although independent genomic responses are associated with parasitism, observed commonalities among these may provide a basis for universally-applicable approaches to targeting parasites through drugs and other interventions. From these genomes, we analysed 0.5 million proteins for their promise as new targets for development of sorely needed drugs, and suggest likely compounds that target these.

## Supplementary Information

Supplementary Information is available in a separate file.

## Acknowledgements

We would like to thank: the WTSI DNA Pipeline teams, particularly Coline Griffiths, Naomi Park, Lesley Shirley, Mike Quail, Dave Willey and Matt Jones; WTSI Pathogen Informatics, especially Jacqueline Keane; Thomas D. Otto for bioinformatics advice; MGI faculty and staff, especially Michael Schmidt, Catrina Fronick, Matt Cordes, Tracie Miner, Robert Fulton and other members of the Project Management, Resource Bank, Library Construction and Data Production teams; Dan Hughes, Matthieu Muffato at the European Bioinformatics Institute, for support running Maker and Ensembl Compara; and Karim Gharbi and his staff at Edinburgh Genomics for support; Verena Gelmedin, Ricardo Fujiwara, Fatima Brazil, the late Purnomo (University of Indonesia, Jakarta), Julie Ahringer, Esther S. Hernández Redondo, Frank Jackson, Elizabeth Redman, Akira Ito, Jenny Saldaña, Marìa Fernanda Dominguez, Wiliam Gause, Mathieu Badets, Irene E. Samonte, Anson Koehler, Martin Nielsen, L. S. Mansfield for sample preparation. The work was funded by Wellcome Trust grant number 206194, US National Institutes of Health (NIH)–National Human Genome Research Institute grant number U54HG003079, National Institute of Allergy and Infectious Diseases grant number AI081803, National Institute of General Medical Sciences grant number GM097435, Medical Research Council (UK) grant number MR/L001020/1. D.W.T., J.E.A., M.L.B., B.M., V.T., Ga.K., Ge.K. and D.R.L. were supported by EU SICA award 242131 ‘Enhanced Protective Immunity Against Filariasis’ (to D.W.T.), a BBSRC/Edinburgh University PhD scholarship and a joint Edinburgh University/James Hutton Institute PhD scholarship (to D.R.L.). J.E.A. was also supported by MRC grant MR/K01207X/1. J.P. and L.S.S. were supported by the National Institutes of Health / NIAID (R21 AI126466) and the Natural Sciences and Engineering Research Council of Canada (RGPIN-2014-06664). R.M.M. is supported by a Wellcome Trust Investigator Award (Ref 106122) and Wellcome Trust core funding to the Wellcome Centre for Molecular Parasitology (Ref 104111). Additional computing resources were provided through Compute Canada by the University of Toronto SciNet HPC Consortium. Schistosome samples were obtained from the SCAN repository (Wellcome Trust grant 104958).

## Author contributions

Full author contributions are detailed in a separate ‘Supplementary Information’ file.

## Author information

Sequence data have been deposited in the European Nucleotide Archive (ENA). Assemblies and annotation are available at WormBase and WormBase-ParaSite (http://parasite.wormbase.org/). All have been submitted to GenBank under BioProjects listed in Supplementary Table 20. The authors declare no competing interests. Correspondence to Matthew Berriman, Makedonka Mitreva and Avril Coghlan.

---

**Extended Data Fig. 1:**
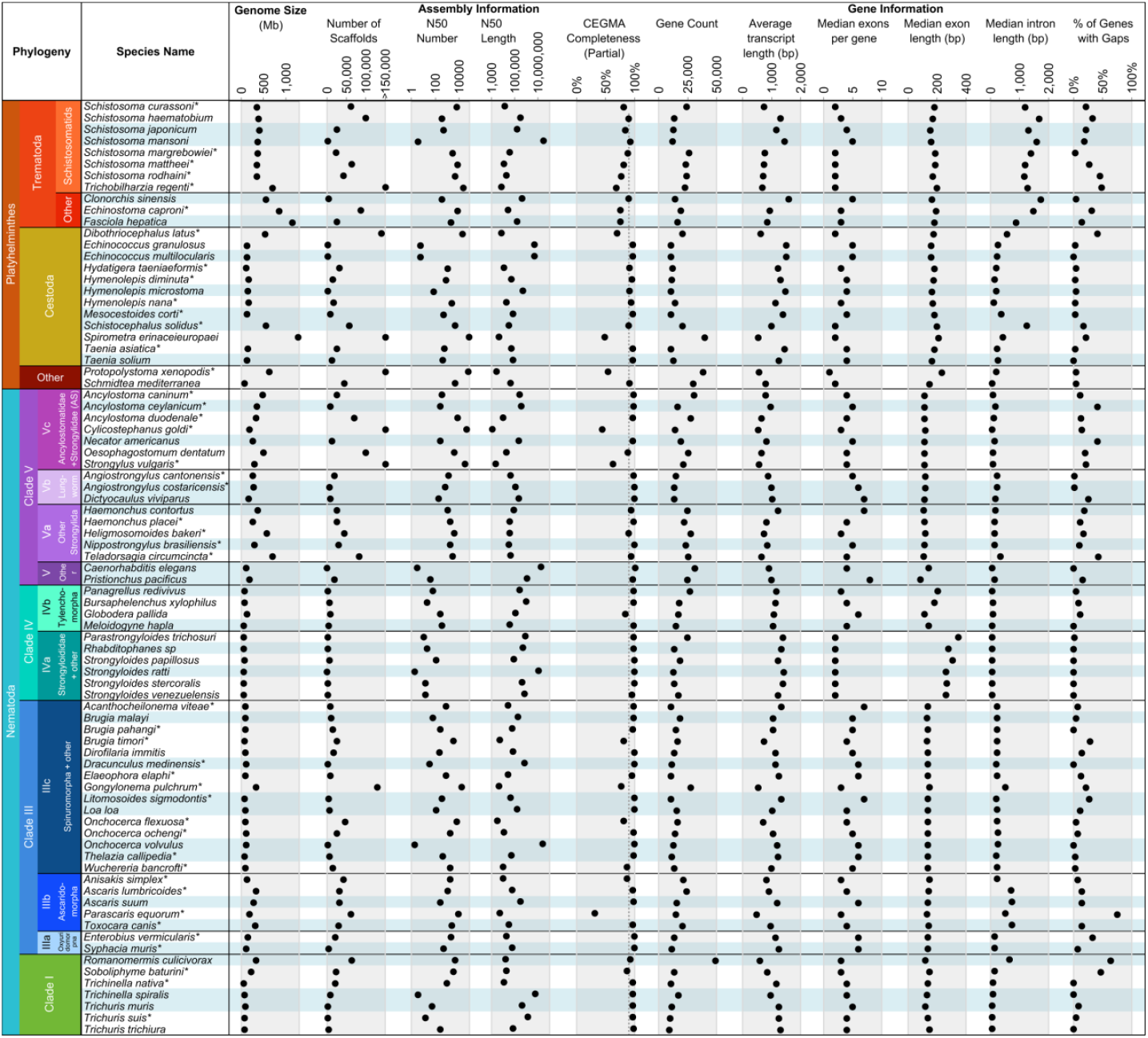
Assembly and gene set statistics. Blue rows indicate the 33 ‘tier 1’ genomes (with high quality assemblies; Supplementary Information: Methods 7). Asterisks indicate the species for which we have sequenced genomes. The ‘% of genes with gaps’ indicates the proportion of predicted genes which contain a sequence gap (i.e., the gene spans multiple contigs).

**Extended Data Fig. 2:**
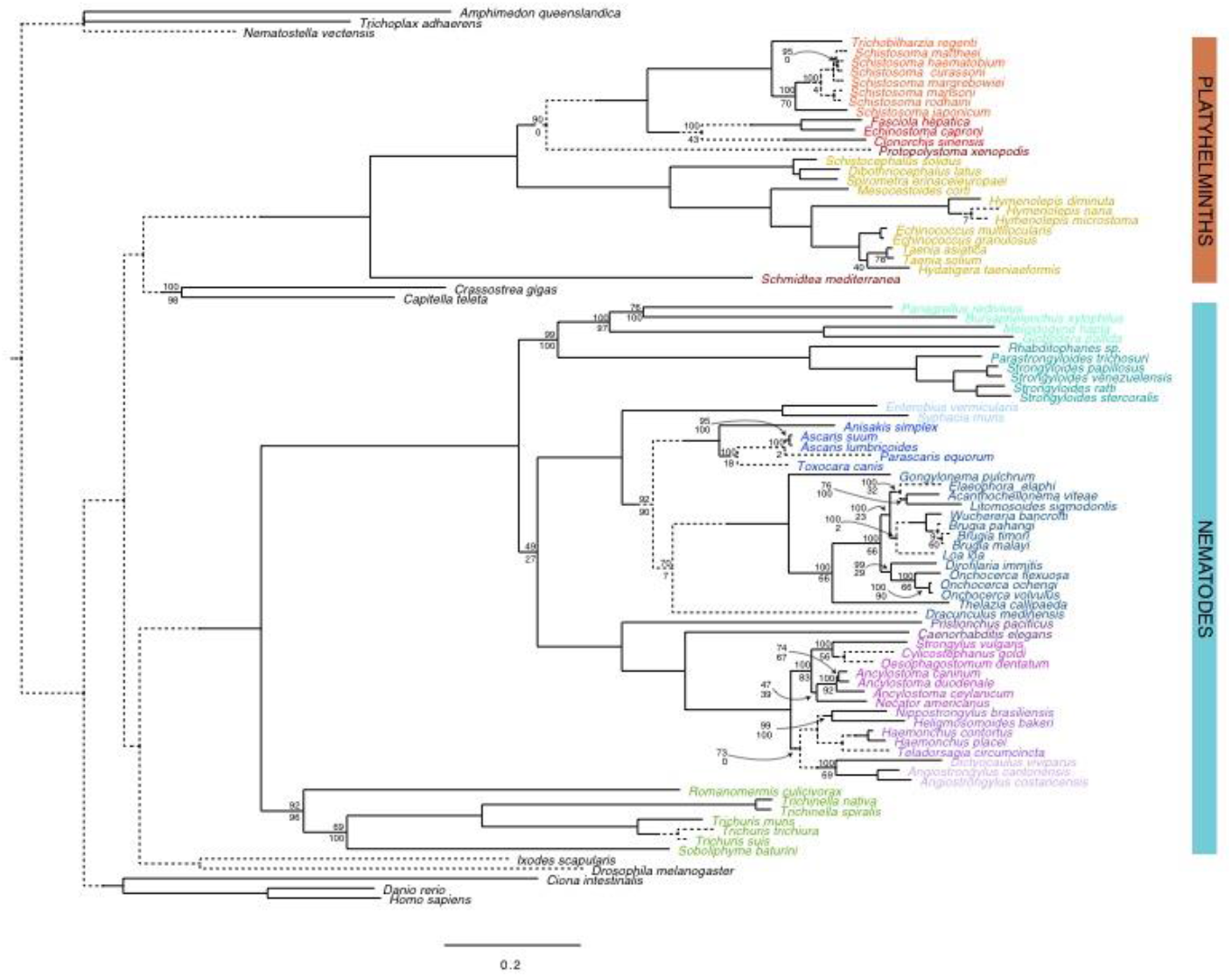
Support for species tree relationships by gene family content. The phylogenetic tree shown is the species tree from single-copy families, also shown in Fig. 1. Here, branches around nodes not supported by gene family content data are shown with broken lines. Values above nodes are bootstrap values from the amino acid matrix analysis; those below nodes are bootstrap values from the gene content data analysis. Where values are not shown next to solid branches, both are 100/100. Where values are not shown next to dotted branches, they are 100 for the amino acid data and zero for the gene content analysis. Contentious issues include the relationships between the monogeneans, cestodes and trematodes within Platyhelminthes; between nematode clades III, IV and V; between Spiruromorpha+other, Ascaridomorpha and Oxyuridomorpha within clade III; and the positions of the genera *Bursaphelenchus* and *Dictyocaulus* within Nematoda (Supplementary Information: Results 2.6).

**Extended Data Fig. 3:**
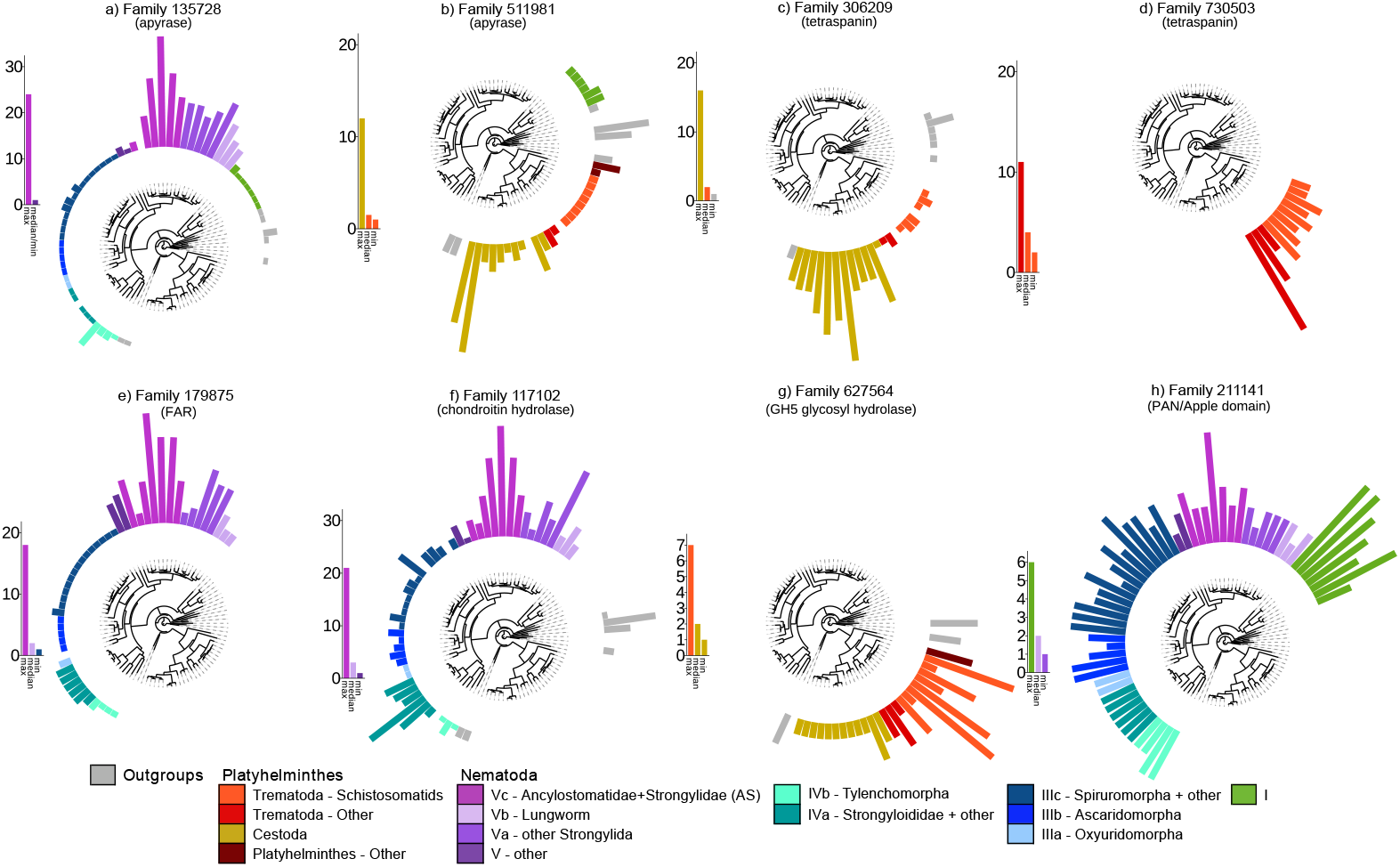
Expanded families involved in parasitism. Gene families with striking variation across species and potential roles in parasitism. Families were defined using Compara. For colour key and species labels, see Fig. 1. The plot for a family shows the gene count in each species, superimposed upon the species tree. A scale bar beside the plot for a family shows the minimum, median, and maximum gene count across the species, for that family.

**Extended Data Fig 4:**
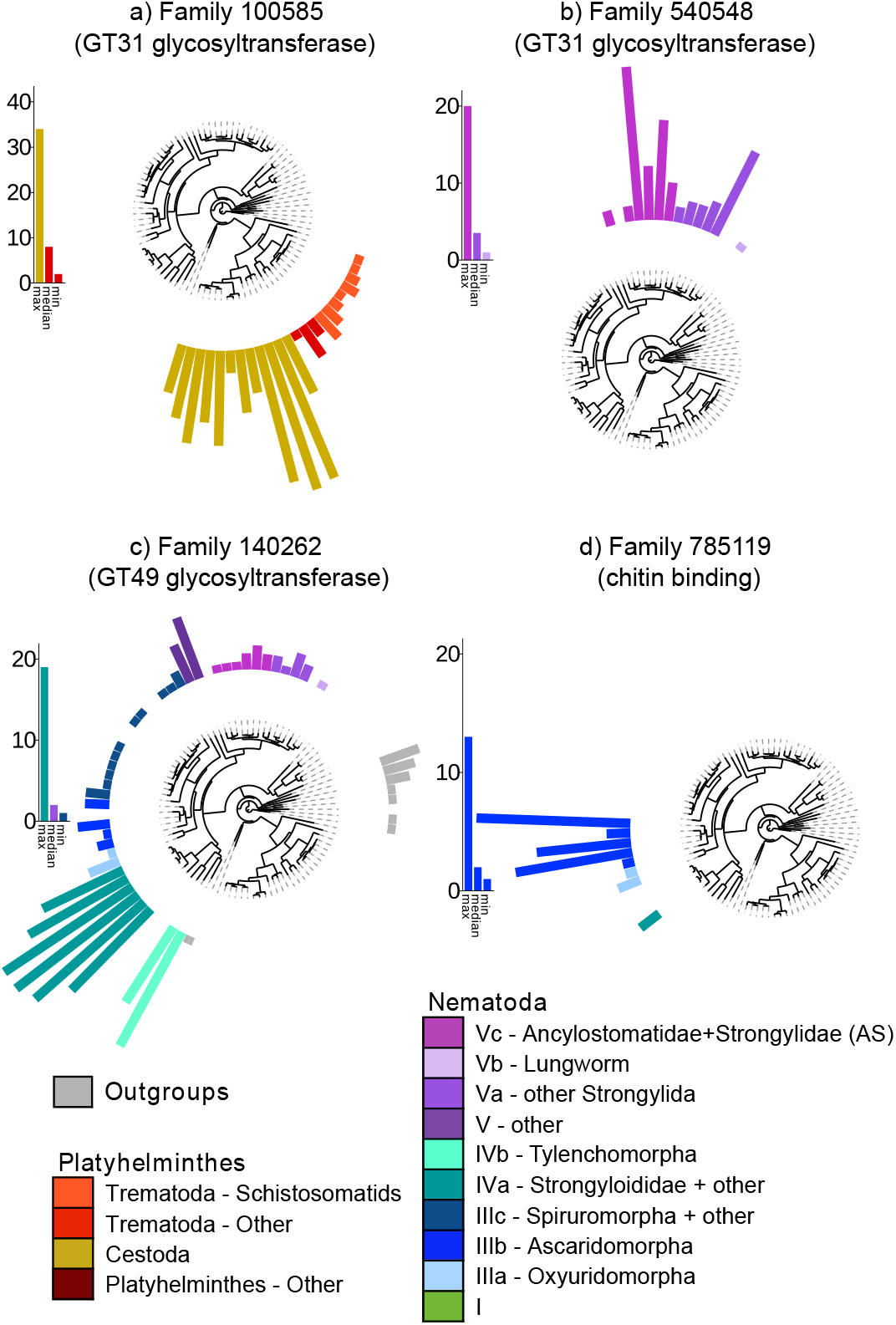
Surface coat expanded families. Expanded families that may be involved in the surface coat. The plot for a family shows the gene count in each species, superimposed upon our species tree. A scale bar beside the plot for each family shows the minimum, median, and maximum gene count across the species, for that family.

**Extended Data Fig. 5:**
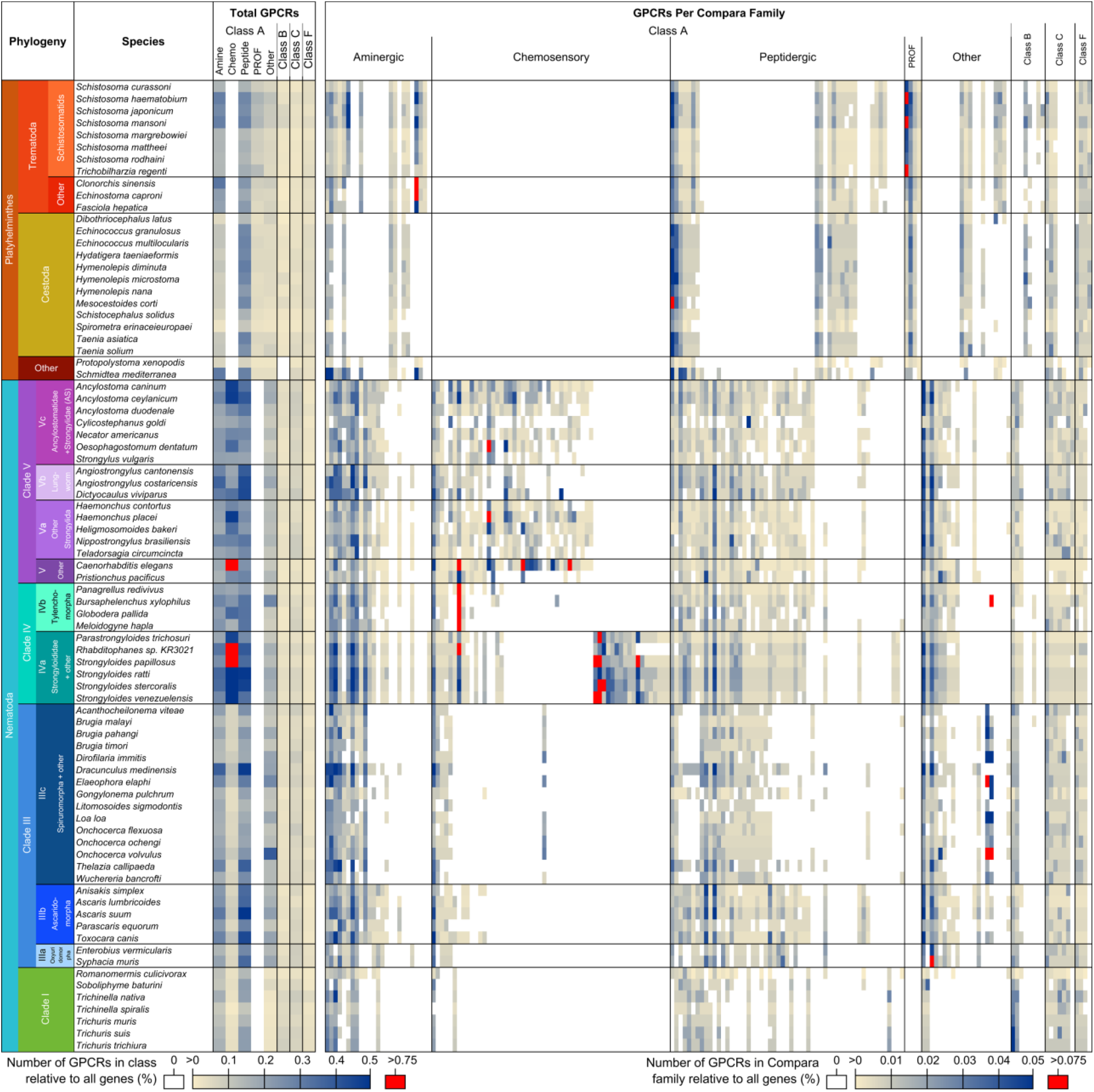
Abundances of GPCRs. Relative abundance profiles for the major categories of GPCRs and the 180 GPCR gene families (columns) represented in at least three platyhelminth or nematode species (Supplementary Information: Methods 19). 50 families present in fewer than three species were omitted from the visualisation.

**Extended Data Fig. 6:**
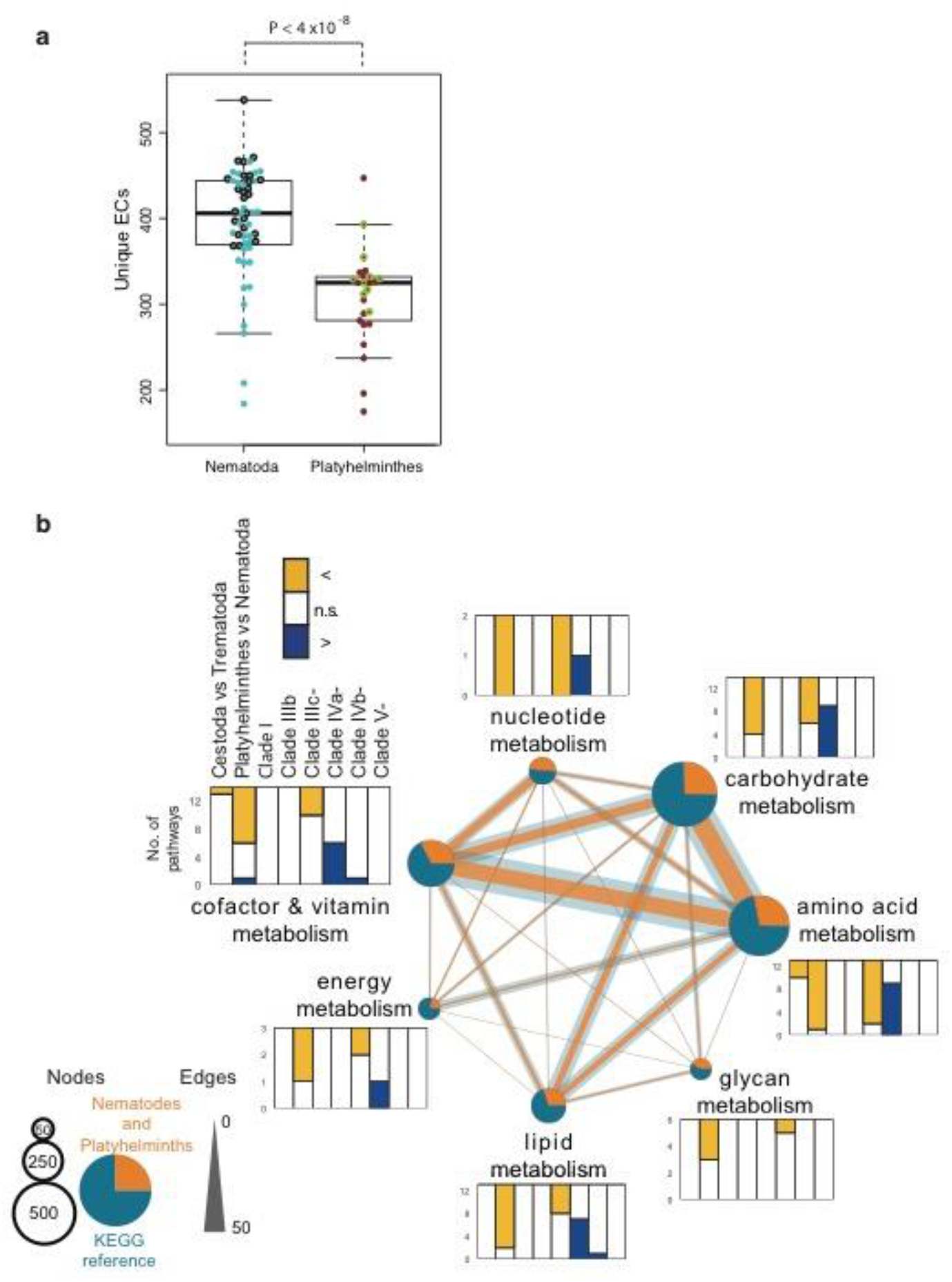
Comparison of EC number annotations and KEGG pathway coverage. (a) Unique enzymes (EC numbers) annotated before using pathway hole-filling. Tier 1 species (with high quality assemblies; Supplementary Information: Methods 7) are highlighted with thick borders. (b) Comparison of KEGG reference pathway coverages (Supplementary Information: Methods 25), using Wilcoxon tests. The pathways are arranged according to superpathways. Nematode parasite groups are compared with the other five nematode sets combined. The total number and overlap in EC numbers of different superpathways are depicted in the central network. The colour of pies and lines indicate the ECs detected in platyhelminths and nematodes, versus the rest of the KEGG reference database. The size of pie charts and lines indicate the amount of EC numbers. The nematode set names are defined as follows: clade I, clade I nematodes; IIIb, clade IIIb nematodes; IIIc-, filaria (=clade IIIb minus 3 non-filaria); IVa-, parasites of clade IVa (i.e. *Rhabditophanes* excluded); IVb-, parasites of clade IVb (i.e. *P. redivivus* excluded); clade V-, parasites of clade V (i.e. *C. elegans* and *P. pacificus* excluded).

**Extended Data Fig. 7:**
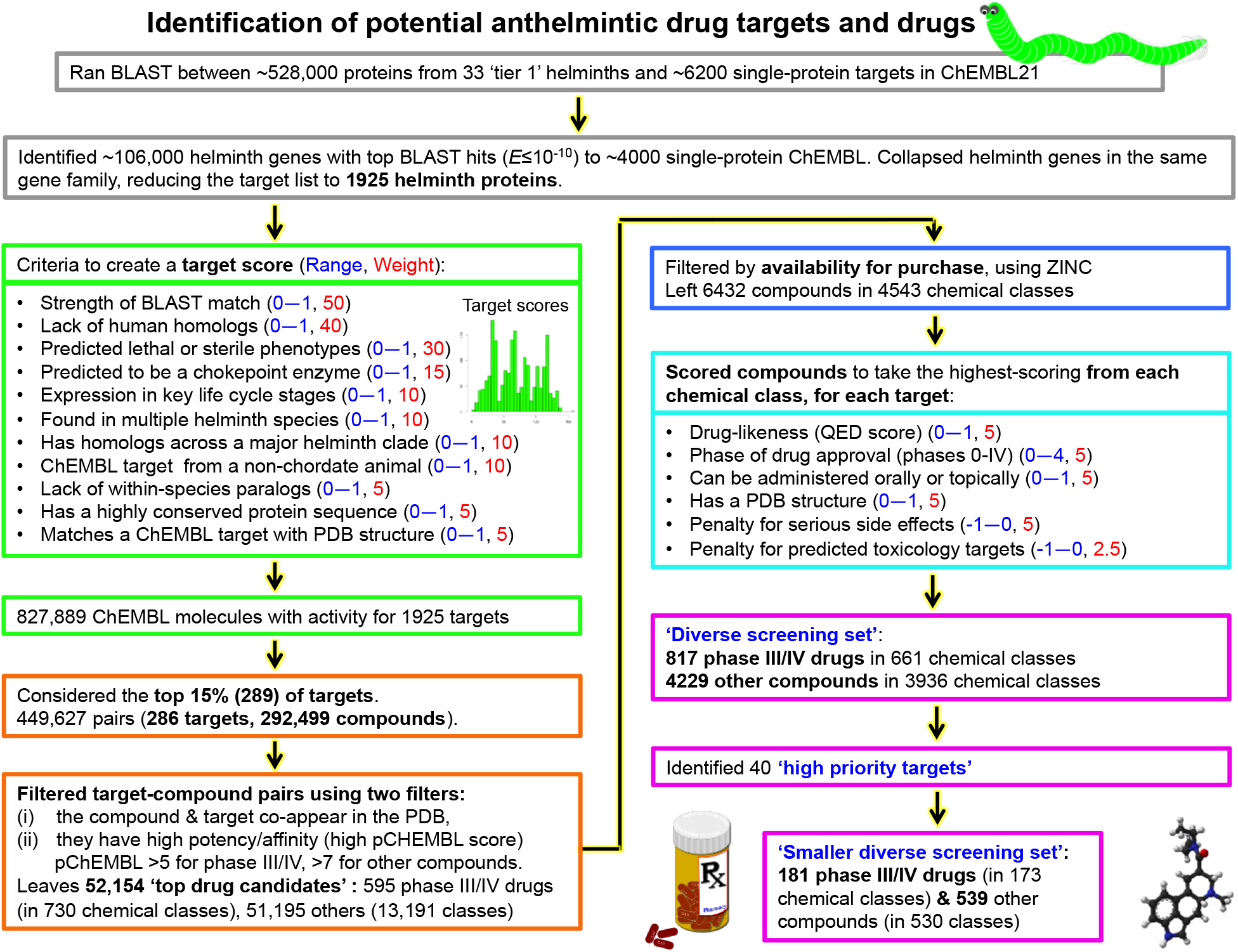
Flowchart of the methods used for identifying potential new anthelmintic targets and drugs using ChEMBL. The following images were used from Openclipart: ‘prescription-bottle-and-pills’ (by algotruneman), ‘famous- and-infamous-molecules-13’ (by Firkin) and ‘worm-gusano’ (by ainara14).

**Extended Data Fig. 8:**
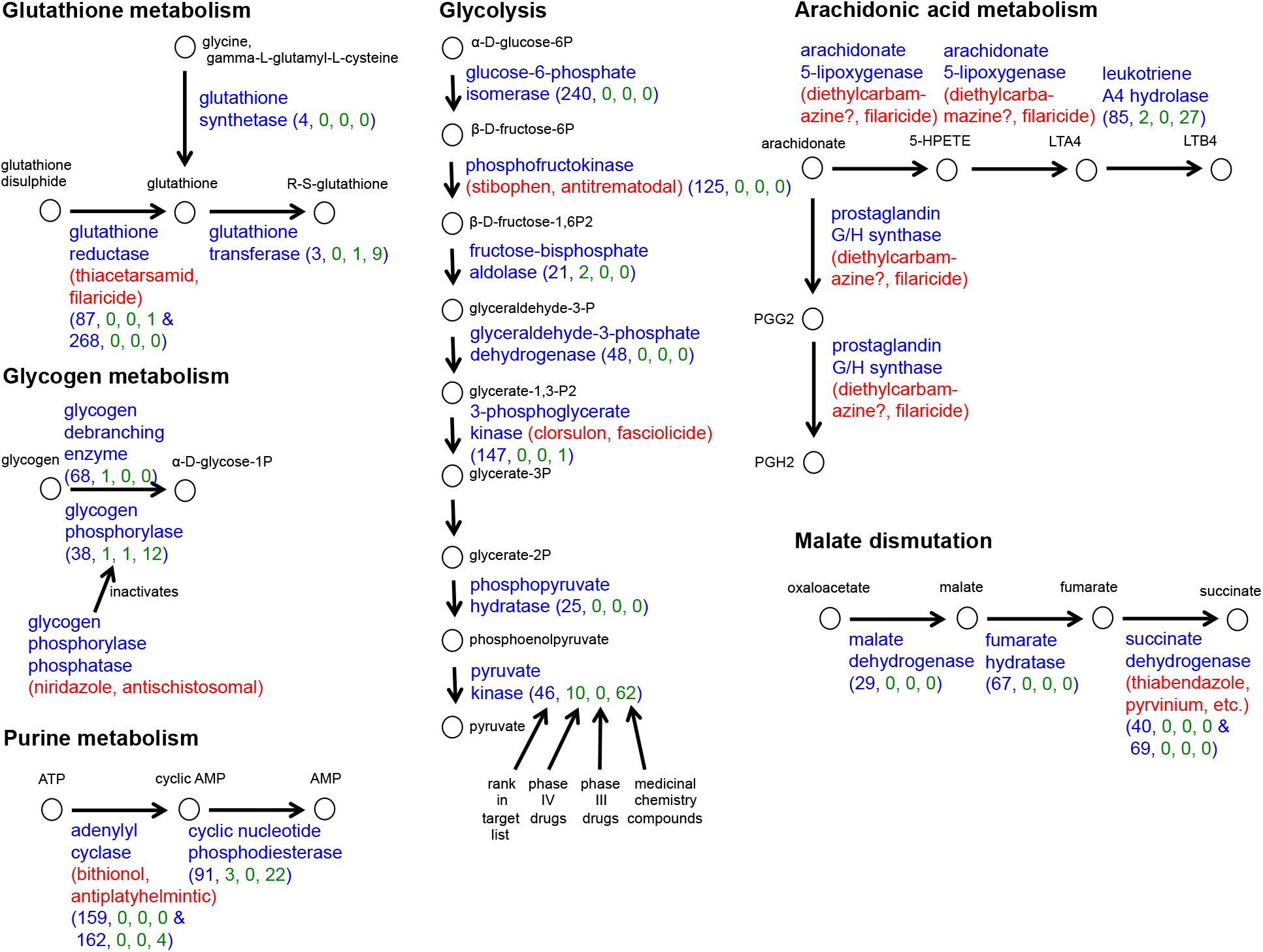
Pathways with known targets. Suggested novel drug targets (from Supplementary Table 21d) that are in the same biochemical pathway as a known or likely anthelmintic drug target (from Supplementary Table 21b). The known anthelmintic drugs for known or likely targets are shown in red. The rank of the suggested target in our target list (Supplementary Table 21d) is given in parentheses, along with the number (in green) of suggested phase III or phase IV drugs and medicinal chemistry compounds (Supplementary Table 21d).

